# Enhanced spine stability and survival lead to increases in dendritic spine density as an early response to local alpha-synuclein overexpression in mouse prefrontal cortex

**DOI:** 10.1101/2023.09.28.559765

**Authors:** Peter J. Bosch, Gemma Kerr, Rachel Cole, Charles A. Warwick, Linder H Wendt, Akash Pradeep, Emma Bagnall, Georgina M. Aldridge

**Affiliations:** Department of Neurology, Carver College of Medicine, University of Iowa, Iowa City; Department of Pharmacology, University of Iowa, Iowa City; Iowa Neuroscience Institute, Department of Neurology, University of Iowa, Iowa City, IA; Institute for Clinical and Translational Science, University of Iowa, Iowa City, IA

**Author notes:** **Conflict of interest:** There are no conflicts of interest. Corresponding Author: Georgina M. Aldridge, 169 Newton Road 36, Pappajohn Biomedical Discovery Building, University of Iowa, Iowa City, 52242.

**Keywords:** Alpha-Synuclein, dendrite, dendritic spine, Parkinson’s disease, Lewy body dementia, neuron, overexpression, pS129, prefrontal cortex, 2-photon

## Abstract

Lewy Body Dementias (LBD), including Parkinson’s disease dementia and Dementia with Lewy Bodies, are characterized by widespread accumulation of intracellular alpha-Synuclein protein deposits in regions beyond the brainstem, including in the cortex. Patients with LBDs develop cognitive changes, including abnormalities in executive function, attention, hallucinations, slowed processing, and cognitive fluctuations. The causes of these non-motor symptoms remain unclear; however, accumulation of alpha-Synuclein aggregates in the cortex and subsequent interference of synaptic and cellular function could contribute to psychiatric and cognitive symptoms. It is unknown how the cortex responds to local pathology in the absence of significant secondary effects of alpha-Synuclein pathology in the brainstem. To investigate this, we employed viral overexpression of human alpha-Synuclein protein targeting the mouse prefrontal cortex (PFC). We then used *in vivo* 2-photon microscopy to image awake head-fixed mice via an implanted chronic cranial window to assess the early consequences of alpha-Synuclein overexpression in the weeks following overexpression. We imaged apical tufts of Layer V pyramidal neurons in the PFC of *Thy1-YFP* transgenic mice at 1-week intervals from 1-2 weeks before and 9 weeks following viral overexpression, allowing analysis of dynamic changes in dendritic spines. We found an increase in the relative dendritic spine density following local overexpression of alpha-Synuclein, beginning at 5 weeks post-injection, and persisting for the remainder of the study. We found that alpha-Synuclein overexpression led to an increased percentage and longevity of newly-persistent spines, without significant changes in the total density of newly formed or eliminated spines. A follow up study utilizing confocal microscopy revealed that the increased spine density is found in cortical cells within the alpha-Synuclein injection site, but negative for alpha-Synuclein phosphorylation at Serine-129, highlighting the potential for effects of dose and local circuits on spine survival. These findings have important implications for the physiological role and early pathological stages of alpha-Synuclein in the cortex.

## Introduction

Lewy Body Dementias (LBD), which include both Parkinson’s disease dementia (PDD) and Dementia with Lewy bodies (DLB) are debilitating multi-system diseases, that result in unpredictable hallucinations, fluctuations in cognition, disturbed sleep, delusions, and depression (Weintraub et al., 2022). Cognitive impairment in PDD/DLB differs dramatically from patients with other forms of dementia; it is characterized by deterioration of executive functions, such as cognitive control, timing, attention, planning, and cognitive flexibility. These characteristics are mediated in part by prefrontal cortical (PFC) circuits (Gnanalingham et al., 1997, Walker et al., 2015, Zhang et al., 2021). Cognitive dysfunction affects 25-30% of Parkinson’s Disease (PD) patients at diagnosis, and cognitive impairments resulting from LBDs have extensive negative impacts on the quality of life, contributing to a loss of independence, disability, and significant caregiver burden (McKeith et al., 2017, Obeso et al., 2017, Zaccai et al., 2005). It is estimated that 80% of PD patients surviving 20 years will develop dementia; as time from onset increases, the chances of developing dementia also increase (Hely et al., 2008, Walker et al., 2015).

Lewy Bodies are large intracellular aggregates that contain the highly expressed protein, alpha-Synuclein (α-Synuclein, α-Syn). Pathologically, PD is characterized by the presence of “Lewy Bodies” in the midbrain, in combination with pallor indicative of loss of dopaminergic neurons in that location (Obeso et al., 2017). By contrast, DLB and PDD cases are often distinguished from PD cases by an increased number of diffuse cortical or limbic aggregates also containing α-Syn, but which can differ in morphology from brainstem Lewy Bodies (McKeith et al., 2017). In both LBD and PD, the majority of α-Syn in aggregates is phosphorylated at Serine-129 (pSer129) and is thought to be an indicator of the main pathological form of the protein (Anderson et al., 2006, Fujiwara et al., 2002, Colom-Cadena et al., 2017).

Evidence for a causative role for α-Syn in PD comes from both human patients and animal models; duplication or triplication of the α-Syn gene, SNCA, leads to familial forms of autosomal-dominant early onset PD and DLB (Spellman, 1962, Singleton et al., 2003). Consistent with the hypothesis that the magnitude of α-Syn expression contributes to disease onset, triplication of SNCA leads to earlier onset and more rapid progression than duplication, which progresses more closely to idiopathic cases (Chartier-Harlin et al., 2004). Efforts to tease apart the role of α-Syn overexpression in animal models have mostly focused on deterioration of motor function. However, it is noteworthy that patients with SNCA triplication are at higher risk for cognitive dysfunction (Chartier-Harlin et al., 2004, Fuchs et al., 2007), whereas some mutations in SNCA lead to parkinsonism with less cognitive effect (Planas-Ballvé and Vilas, 2021).

Although cell loss (including dopaminergic and cholinergic projection neurons) is seen in patients with α-Syn aggregates in the brainstem, and undoubtably contributes to symptoms characteristic of LBD, there are examples of individuals with pathology in the cortex with less severe pathology in the brainstem (Zaccai et al., 2008). Furthermore, understanding the implication of cortical α-Syn aggregates in the absence of severe dopaminergic deficits may aid our understanding of heterogeneity in PD/PDD/DLB. In particular, the associative cortices, including PFC, often show α-Syn pathology in patients with DLB and PDD. The PFC is central to many higher order brain processes in humans, including attention, habit formation, decision making, working memory, planning, reward prediction, and emotional and inhibitory control (Ohnuki et al., 2021, Bittar and Labonté, 2021). Both DLB and PDD patients have a pattern of cognition that include deficits in executive functions (Aldridge et al., 2018, Smirnov et al., 2020). In addition to α-Syn pathology, there is evidence from a small number of LBD autopsies that the PFC has decreased density of dendritic spines (Kramer and Schulz-Schaeffer, 2007), although co-morbid mixed pathology, such as beta-amyloid, was not clearly ruled out and alternative methods do not support a loss of synapses in pure Lewy Body Dementia (Hansen et al., 1998). Alternative mechanisms for how local aggregation of α-Syn might impact cortex include pre- and post-synaptic protein deficits (Kramer and Schulz-Schaeffer, 2007) and other circuit related deficits (Schulz-Schaeffer, 2015, Nikolaus et al., 2009), which could have a profound influence on behaviors and cognitive functions mediated by this region. However, it is unclear if α-Syn pathology directly or indirectly causes circuit level dysfunction in PFC cells.

Dendritic spines are small protrusions that extend from neuronal dendritic shafts and are the post-synaptic site for most excitatory neurons in the brain (Hedrick et al., 2022). Dendritic spine populations exhibit experience-driven plasticity, which manifest as dynamic changes in their anatomy (size, morphology, and length) and numbers (Nava et al., 2017, Pan et al., 2010). Transgenic overexpression of human α-Syn driven by the platelet-derived growth factor (PDGF-b) promoter in mouse (Blumenstock et al., 2017) and α-Syn preformed fibril injections into dorsal striatum after 5 months were associated with decreases in spine density in Layers I and Layer IV/V cortical neurons (Blumenstock et al., 2017). However, these studies did not examine the effect of α-Syn overexpression isolated to a cortical region. Further, single time representations of dendritic spine anatomy and number, while useful, miss the rapid temporal dynamics that occur on the days-to-weeks’ timescale.

In our previous work, we found an increased dendritic spine density in the medial PFC following local α-Syn overexpression, using a single post-mortem timepoint paired with Golgi staining (Wagner et al., 2020). Our data suggested that 10 weeks following injection, mice with α-Syn overexpression had greater local spine density than mice injected with a control virus. Here, we used longitudinal 2-photon transcranial imaging to identify transient spine changes and determine the cause of this increase in spine density by examining Layer V neuronal tufts (visualized in Layer I) on a weekly basis before and after local viral overexpression of human α-Syn in the mouse PFC. Our results provide new data to show that α-Syn overexpression initially increases local dendritic spine density beginning 5-weeks post-AAV injection, and this increase is secondary to increased new-spine stability and survival in the region of overexpression.

## Results

### Alpha-Synuclein overexpression dynamically increases dendritic spine density in apical tufts of Layer V cells in the prefrontal cortex

Using the *Thy1-YFP* transgenic mouse line and AAV-mediated overexpression of human α-Syn in the PFC, we studied the dynamic changes in dendritic spines in awake, head-fixed mice using 2-photon microscopy. *Thy1-YFP* mice (Feng et al., 2000) express yellow fluorescent protein (YFP) in a subset of Layer V (and a smaller number of Layer II/II) neurons in the cerebral cortex, which fill the whole cell with YFP, including neurites and dendritic spines. We therefore used this mouse line to perform repeated imaging of dendrites and their dendritic spine protrusions. Our strategy combined cranial windows with intracranial injections through a pre-drilled hole, which had been plugged with silicon to allow for injection needle entry (**Fig. 1A, B**). Both groups received unilateral injections of AAV. Control animals received a virus encoding for the mCherry fluorescent protein. α-Syn overexpression (OE) mice received a mixed combination of the AAV coding for overexpression (OE) of human α-Syn and that for mCherry (henceforth α-Syn OE). We targeted the PFC, mainly encompassing the M2 (Secondary motor), Cg1 (Cingulate) and PrL (Pre-limbic) regions; some overexpression also extended into the M1 region (**Fig. 1C, D**). Immunofluorescence staining using an antibody targeted to phosphorylated α-Syn at Serine 129 (pSer129) and confocal microscopy demonstrated robustly expressed pSer129 throughout transduced cells, with strong expression in the soma and nuclei of neurons. There was substantial staining in the dendrites and some spines also displayed positive pSer129 staining at 70 days post injection (**Fig. 1E-G**).

**Figure 1.**
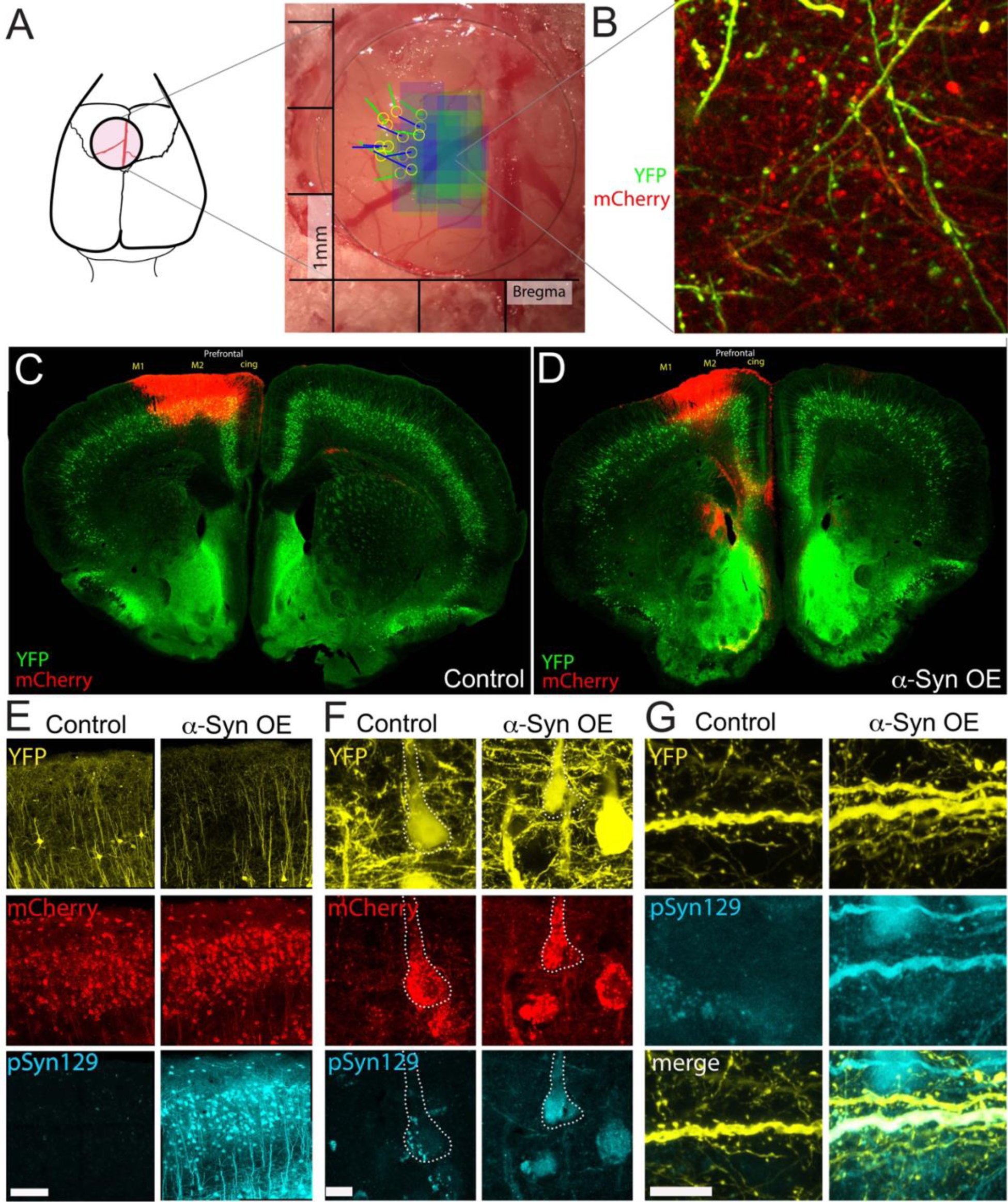
Experimental design for overexpression of human alphα-Synuclein in the mouse PFC. (A) We placed round cranial windows over the putative imaging site. AAV injections were performed after two imaging sessions, through a pre-drilled hole plugged with silicon. Injection sites for mice in the study are shown as yellow circles, with the lines (blue = control, green = α-Syn OE) representing the angle of injection and the colored boxes (blue = control, green = α-Syn OE) representing the imaging sites of apical tufts. (B) Representative image from a highly transduced area. (C-D) Example of the localized spread of transduction in a *Thy1-YFP* coronal section, showing AAV mCherry expression in a control (C) and α-Syn OE (D) section, localized to the mPFC (M2, cingulate) region. (E) Overexpression of human-α-Syn leads to robust phosphorylation at pSer129 in treated mice, including in soma (F) and dendrites (G). Scale bar in (E) 100 mm. Scale bar in (F) and (G) 10 mm.

Mice were implanted with cranial windows and a secure headplate, which stabilized the head whilst the mouse could run on a floating platform during imaging to minimize stress. This strategy provided stable imaging and sufficient consistency to image the same dendrites each week. We placed a cranial window 3 weeks prior to 2-photon transcranial imaging onset and imaged weekly for up to 11 weeks (**Fig. 2A**). Data collection was ended (or weeks were excluded) when imaging quality was not sufficient to assure quantification of smaller spines, as determined by blinded investigators. Mice (age at imaging start, Control: 38.9 ^+^/-5.3 weeks, n=6; α-Syn OE: 39.9 ^+^/-4.4 weeks, n=8, mean+/-SEM) were initially imaged for 2 weeks (Week −1 and Week 0, “PRE”) to establish the pool of examinable dendrites and then injected with the relevant viruses (**Fig. 2A**, **Fig. 1A**). We then continued imaging the same dendrites for a further 9 weeks (“POST”, **Fig. 2A**).

**Figure 2.**
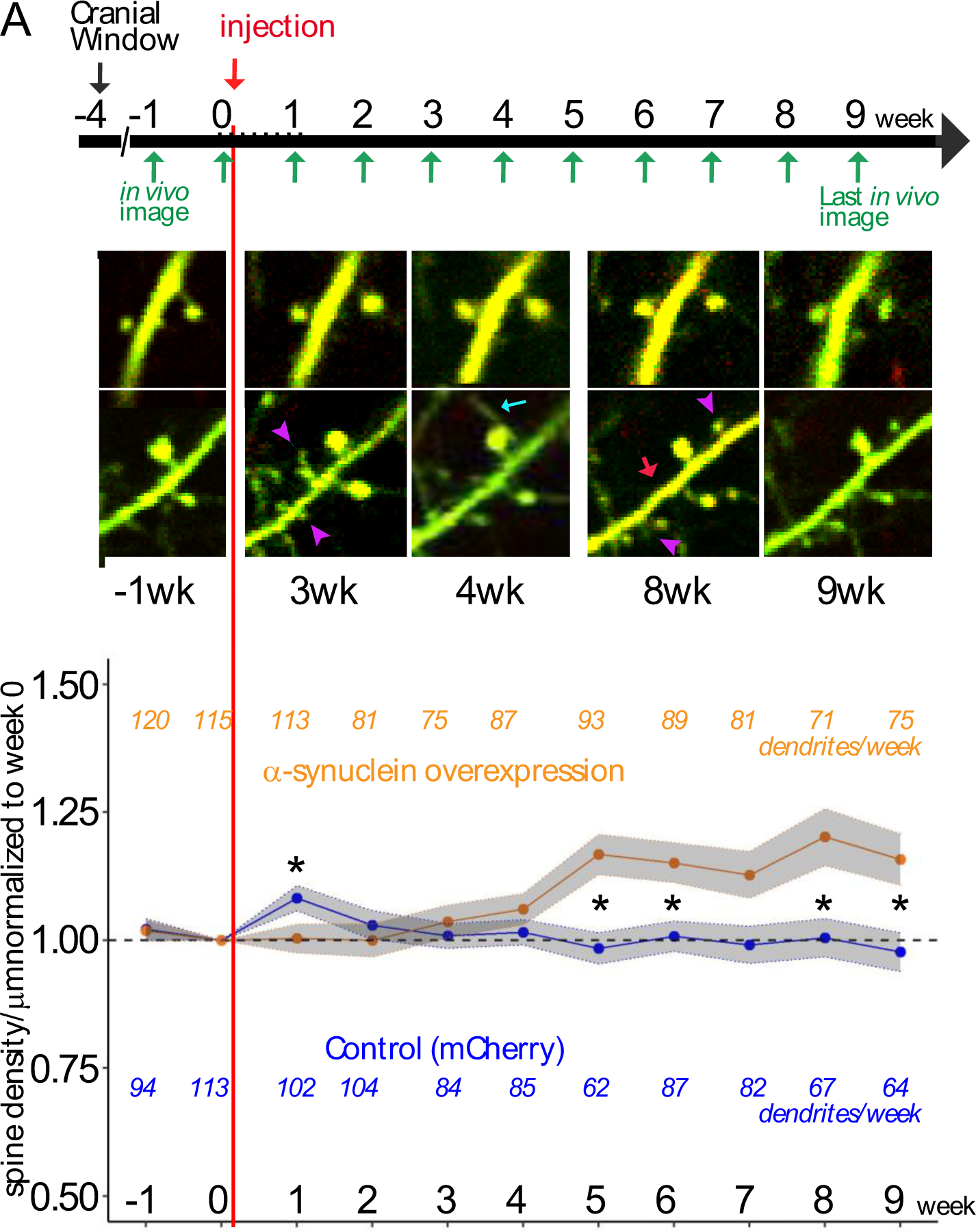
Alphα-Synuclein overexpression causes increases in dendritic spine density on individual dendrites. (A) 2-photon imaging was performed 3 weeks after cranial window surgery to generate 2 weeks of pre-treatment imaging of dendrites (week −1 and week 0, “PRE”), followed by AAV injection of control or α-Syn OE constructs immediately after the week 0 imaging session. Nine weeks of 2-photon microscopy was performed (“POST”) on the same pool of dendrites that were established in weeks −1 and 0. (B) Representative, single plane images of control (top) and α-Syn OE (bottom) dendrites. Purple arrowheads represent new spines; cyan arrow is a new filopodia; red arrow is a lost spine, each identified by comparison of the full 3D stacks, not visible here. (C) Dendritic spine density increases in α-Syn OE animals compared with control. Each dendrite was divided by its pre-treatment density to evaluate the effect on individual dendrites. The orange and blue colored numbers represent the number of dendrites measured for each group at each week; dendrites were excluded at some weeks due to changes in the cranial window transparency or angle, equipment malfunction, and the covid pandemic. α-Syn OE led to increased dendritic spine density starting at week 5 and continuing for the duration of imaging (*p < 0.05).

We found that spine density in mice overexpressing human α-Syn was lower than control animals in the week immediately following injection, but then significantly increased in the apical tufts of Layer V PFC neurons on a weekly basis, while spine density significantly decreased on a weekly basis in dendrites from mice injected with control virus (**Fig. 2B, C, Supp Fig 1.;** Control vs. α-Syn p=0.006, week*treatment p<0.001, linear mixed model). α-Syn OE mice had significantly higher spine density at week 5, 6, 8 and 9 post-AAV expression, proportional to Week 0, via weekly Control vs. α-Syn comparisons. We also noted an increase in control dendritic spines one-week post-injection, potentially in response to the viral tranduction. However, this effect was no longer evident by Week 2 (**Fig. 2C**).

### Alpha-Synuclein overexpression in the PFC leads to increased persistence and survival of new spines, with no significant change in formation or elimination rates

The increased spine density we observed began at 5 weeks post injection and continued for the duration of the study (**Fig. 2C**). We therefore asked whether this increased spine density was due to alterations in generation of new spines, changes in elimination, or differential spine persistence. First, we investigated spine formation and elimination densities. To compensate for the infrequency of these events, formation and elimination densities per week were grouped into early (week 1-3), mid (week 4-6) and late (week 7-9). This analysis revealed no significant differences in spine formation (Two-Way ANOVA, Treatment F(1,456)=1.634, p=0.2018, **Fig. 3A**) or spine elimination (Two-Way ANOVA, Treatment F(1,473)=2.039, p=0.1540, **Fig. 3B**) between the groups. Next, we sought to determine if the increased density was therefore due to increased persistence of newly-formed spines. Newly formed spines can be classified by whether they remain the week after they form (persistent) or recede (transient), so we identified what percentage of newly formed spines were still there the week after appearing (new-persistent) and whether there were any differences in persistence between control and α-Syn OE groups following injection (**Fig. 3C**). It was not possible to identify persistence rates prior to injection given the limited number of imaging weeks.

**Figure 3.**
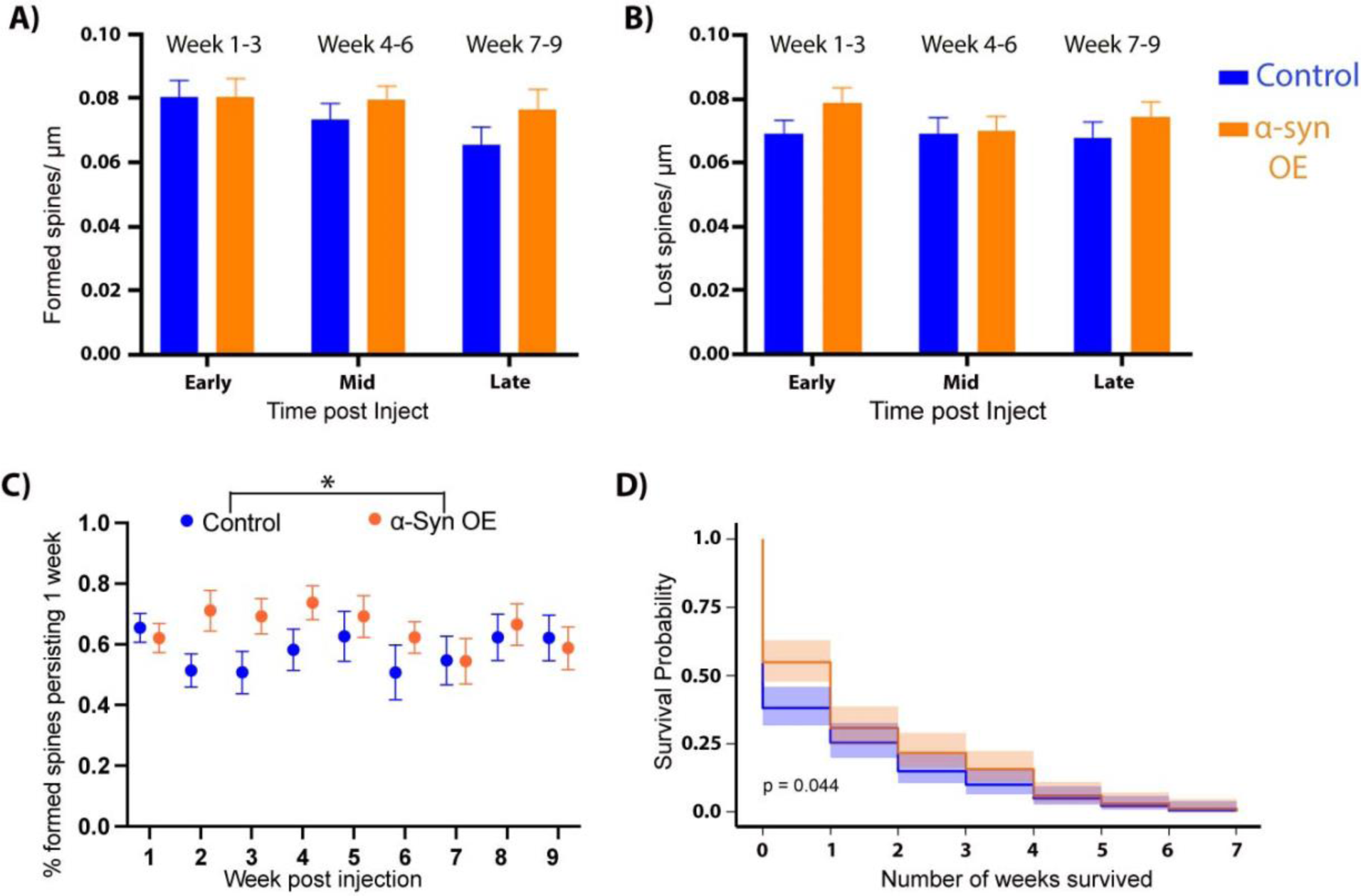
Increases in spine density following α-Syn overexpression are due to increased spine persistence. The increase in spine density could not be attributed to significant differences in Spine Formation. **(A)**: 2-way ANOVA, F(1,456)=1.634, p=0.2018), or **Spine Elimination (B):** 2-way ANOVA, F(1,473)=2.039, p=0.1540**. Persistence (C):** The percent of new spines, grouped by dendrite, that persisted for at least one week following their appearance was higher in the α-Syn OE mice (Odds ratios (OR) for spine survival by TREATMENT (α-syn vs. control): OR: 1.30 CI: 1.01-1.67, p =0.040) compared with controls, with no significant main effect of time. **Survival (D):** Spines that first appeared in weeks 1, 2, or 3 post-injection survived significantly longer in the α-Syn OE mice compared with spines in the control-injected mice (p=0.044), Kaplan-Meier curve.

This analysis demonstrated that dendrites from mice with local α-Syn OE had a significantly higher percentage of new-persistent spines in the weeks after AAV injection (Odds ratios (OR) for spine survival by TREATMENT (syn vs. control): OR: 1.30 CI: 1.01-1.67, p =0.040, and WEEK: 0.98, CI: 0.94-1.02, p = 0.3). An interaction between time and TREATMENT was examined and not found to be significant (**Fig. 3C**). We further investigated the survival of newly formed spines, so we generated survival curves for each new spine identified during the first 3 weeks after the treatment AAV injections had been performed. We found that dendritic spines in the α-Syn OE group survived for a significantly longer time that controls (Kaplan-Meier, p=0.044, **Fig. 3D**). Overall, these data suggest a mechanism whereby an increase in spine survival and persistence leads to a relative increase in spine density by 5 weeks after α-Syn OE in the PFC. Due to limitations of imaging in aged YFP-expressing mice, which have increased background fluorescence, morphological classifications were unreliable between blinded observers; thus, we excluded morphological analysis.

### AAV expression of mCherry inhibits genetically expressed YFP fluorescence

While characterizing dendrites in vivo to determine whether they contained mCherry expression (using laser excitation at 1040 nm), we noted a potential confound. We found that dendrites with high levels of mCherry lost visualization or expression of YFP (Supplemental Fig 2). Dendrites that demonstrated this loss of YFP over time were excluded from all analysis. Thus, in vivo experiments described above are limited to dendrites within the transduction region (affected by the local microenvironment) but excludes dendrites with the highest potential transduction. The few dendrites that were positive for mCherry in vivo, but which didn’t show YFP loss were graphed separately for illustration (Supplemental Fig 1). The finding that genetically encoded Thy-YFP is suppressed in cells with high levels of mCherry is important for design of future studies. We have not noticed this phenomenon with expression of α-Syn alone (not shown) and there is very little interference when mCherry expression is reduced using a preceding internal ribosome entry site (IRES, see confocal study below), but this lower expression level cannot be visualized well by 2-photon. Importantly for evaluating conclusions of this study, mCherry was used in both the treatment group (AAV-α-Syn mixed with AAV-mCherry) and controls (mCherry-only AAV). mCherry positive dendrites and spines did not display evidence of degeneration; however, we did not attempt quantification of spine density using the red channel, as smaller spines could not be accurately compared once the YFP faded. To more accurately address the effect of local α-Syn, we also qualitatively scored the relative amount of transduction (passing axons and dendrites) at each micro-imaging location as an indicator of the relative exposure to localized expressed protein (Supplemental Fig 1C + D). These graphs are included in supplemental for illustrative purposes (given the low sample size in some sub-groups).

### Increased dendritic spine density in the PFC after local alpha-Synuclein overexpression is primarily driven by cells negative for pSer129 expression

There were a small subset of dendrites in the 2-photon dataset that could positively be identified during in vivo imaging as being transduced via mCherry expression (**Supplemental Fig 1A + B**). However, most of the dendrites measured were mCherry-negative by 2-photon imaging, thus likely non-transduced or transduced with lower-copy levels (as live 2-photon imaging is not as sensitive as confocal microscopy on fixed sections). To assess direct vs. indirect effects of α-Syn OE on spine density, we designed an experiment to more accurately measure spine density based on cells that contain phosphorylation of α-Syn at Ser-129 (pSer129). Because the *Thy1-YFP* line contains high numbers of labeled neurons in older adult animals in Layer V of the cortex, we utilized the *Thy1-GFP* line (Feng et al., 2000), which contains fewer labelled cells and which would allow us to test this hypothesis. We bilaterally injected virus into the PFC of *Thy1-GFP* mice: the mCherry control virus was injected into one hemisphere and a human-α-Syn-IRES-mCherry construct was injected into the contralateral hemisphere of the same mouse. We collected brains 6 weeks after injection and immunostained for GFP and pSer129 to identify pSer129 positive and negative cells from the α-Syn OE hemisphere and compare them with control cells from the contralateral hemisphere. As with *Thy-YFP* animals, we again found GFP fluorescence was reduced, in this case specifically when mCherry alone was overexpressed under the control of the CAG promoter. This prevented accurate spine density of mCherry-positive cells on the mCherry side, even when cells were stained using antibodies to GFP. By contrast, GFP-signal drop did not occur to the same extent when mCherry was expressed using a virus that containing “α-Syn-IRES-mCherry” (injected on the α-Syn OE side). We were therefore able to assess dendritic spine density of basilar dendrites of layer V pyramidal neurons to compare 3 groups: 1) control hemisphere/mCherry-negative; 2) α-Syn OE hemisphere/pSer129-positive; and 3) α-Syn OE hemisphere/pSer129-negative (**Fig. 4A**). After controlling for distance from soma using bins of 0-30 µm, 30-75 µm, and 75+ µm, we found a significant effect of Bin (F(2,250)=21.83, p<0.0001) and Treatment (F(2,250)=4.147, p=0.0169) using 2-way repeated measures ANOVA (**Fig. 4B, C**). Tukey’s multiple comparisons test revealed significant differences in the 75+ µm bin between the α-Syn OE hemisphere/pSer129-negative cells and the other two groups: control hemisphere/mCherry-negative (p = 0.0077, Tukey’s post-hoc test); and α-Syn OE hemisphere/pSer129-positive (p = 0.004, Tukey’s post-hoc test). There was no significant difference in the 75+ µm bin between control hemisphere/mCherry-negative and α-Syn OE hemisphere/pSer129-positive (p = 0.6413, Tukey’s post-hoc test). Our findings reveal that increased spine density early after localized α-Syn OE in the mouse PFC is likely driven by pSer129-negative neurons in the region of transduction, while those expressing pSer129 show no clear differences in spine density (**Fig. 4B, C**), in line with the overall results from our 2-photon experiments.

**Figure 4.**
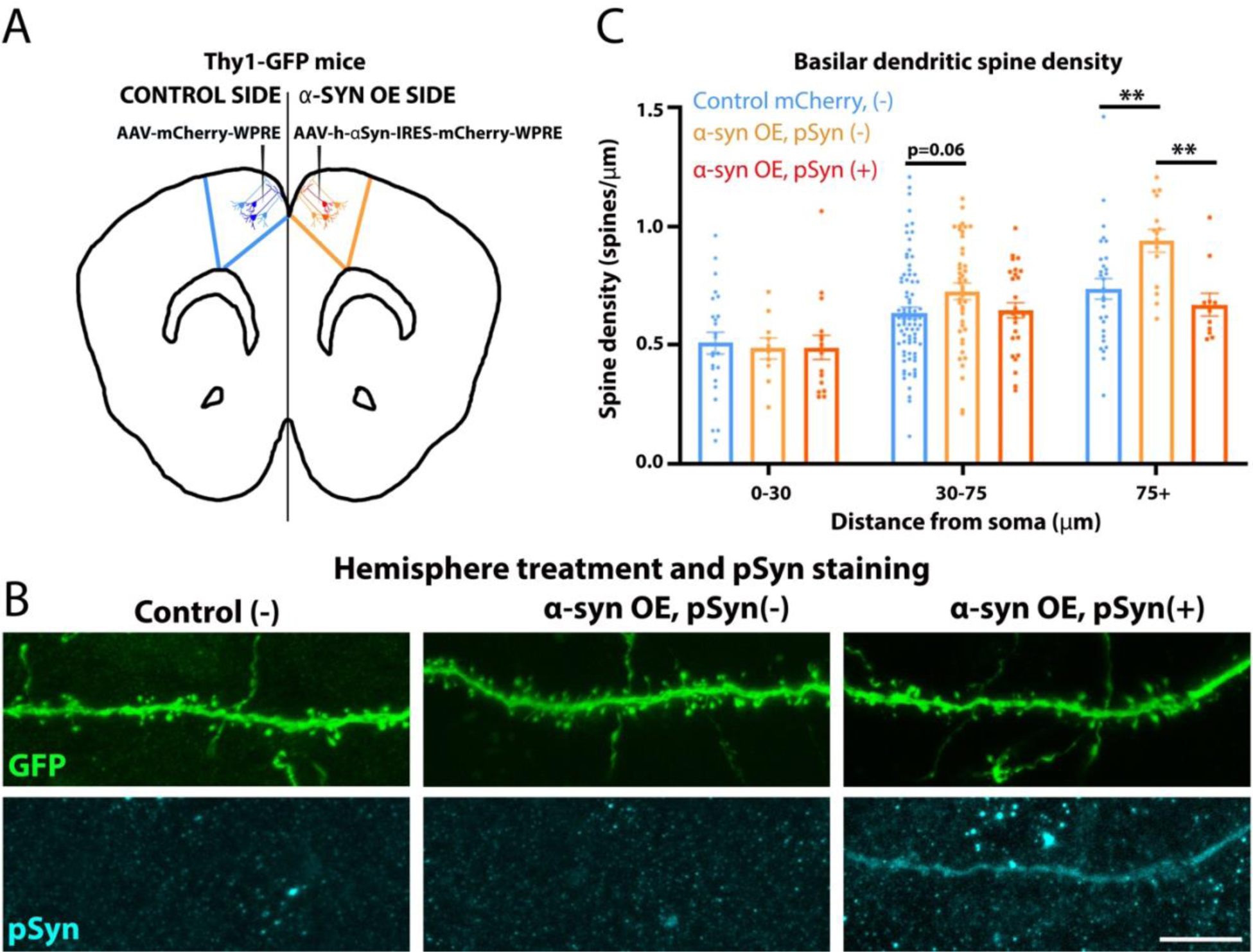
Increased dendritic spine density in response to alphα-Synuclein overexpression occurs in pSer129-negative neurons within the injection location. (A) *Thy1-GFP* mice were injected bilaterally into the PFC: control (AAV-mCherry) in one hemisphere and α-Syn OE (AAV-α-Syn-IRES-mCherry) in the contralateral hemisphere. (B) Confocal imaging of basilar dendrites of *Thy1-GFP* mice, perfused 6 weeks post-injection. Sections were immunostained with antibodies against GFP and pSer129-α-Syn and dendritic spines were counted. (C) Basilar dendrites were analyzed in bins from 0-30, 30-75 and 75+ mm from the soma and grouped according to the hemisphere treatment (control or α-Syn OE). For the α-Syn OE hemisphere, cells were further divided into pSer129-α-Syn positive or negative. The pSer129-α-Syn negative dendrites had greater spine density than controls and pSer129-α-Syn positive dendrites (2-way ANOVA, F(2,250)=4.147, p=0.0169)), which was significantly different at 75+ mm from the soma via Tukey’s post-hoc tests (Control-negative vs. α-Syn-pSer129 negative p = 0.0077; α-Syn-pSer129 positive vs. α-Syn-pSer129 negative p = 0.004).

## Discussion

α-Syn pathology in the cortex is a defining pathological feature of dementia in synucleinopathies, and yet it is unclear how pathology in this region impacts function. Since α-Syn is a synaptic protein, we hypothesized local overexpression would alter dynamics of the synapse and local circuits, leading to alterations in dendritic spines. Turnover of dendritic spines in Layer V cortical pyramidal neurons is an essential functional process, allowing plasticity of these neurons within relevant circuitry (Gemin et al., 2021, Rajkovich et al., 2017). To isolate the effect of α-Syn overexpression on cortical cell function, we combined cranial windows with awake, longitudinal 2-photon imaging to track dendritic spine density before and 9 weeks following human-α-Syn OE in the mouse PFC. Our results suggest that early α-Syn OE targeted locally to the cortex leads to dynamic increases in spine density driven by increased survival of new-persistent spines, rather than significant changes in spine formation or elimination. Next, by using post-mortem confocal microscopy comparing the hemispheres of mice unilaterally overexpressing α-Syn, we were able to show that spines within the injection site, but negative for pSer129-α-Syn phosphorylation had the highest spine density compared with both neurons positive for pSer129-α-Syn and neurons in the contralateral (mCherry virus injection) location. Interestingly, we did not detect any spine density differences in pSer129-positive neurons within the injection site compared with the control side, suggesting that this marker of pathological phosphorylation alone, over this time frame, is not sufficient to cause spine loss. Our results have implications for understanding the health of neurons within the local circuit of neurons developing phosphorylated α-Syn. Overall, these findings are consistent with our previous study in post-mortem mice and provide new insights into the etiology of this early response to overexpression.

There is some evidence that changes at the synapse precede neuronal cell loss that is characteristic of PD and LBD (Calo et al., 2016, Tanji et al., 2010, Kramer and Schulz-Schaeffer, 2007); therefore, understanding early synaptic changes will likely prove beneficial to understanding disease progression as well as the role of native α-Syn. Previous work has investigated α-Syn perturbations in rodents via expression level changes and mutations, as well as in multiple age groups and brain regions relevant to PD and LBD. Studies of the effect of synuclein models on dendritic spine encapsulate brain regions relevant to these diseases; including olfactory bulb (Neuner et al., 2014), hippocampus (Winner et al., 2012), striatum (Graves and Surmeier, 2019) and cortex (Blumenstock et al., 2017, Wagner et al., 2020). Independent analysis of individual regions allows for isolated modelling of aspects of PD/LBD biology to explore differential cell vulnerability, as heterogeneity in pathological distribution and symptoms are a key feature of synucleinopathies (Carceles-Cordon et al., 2023). Our results suggest that our early local cortical overexpression model displays some hallmarks of synuclein pathology (e.g., phosphorylation at Ser129 (Colom-Cadena et al., 2017)) and not others (staining is homogeneous, rather than aggregated, **Fig. 1F**).

Our previous study suggested that local increases of α-Syn in the PFC led to higher dendritic spine density at 10 weeks following viral overexpression using Golgi staining (Wagner et al., 2020). However, it was limited by a single timepoint. The results of our current work support the conclusions of our previous study and build on them by establishing a time-course, which shows that spine density changes can occur much sooner in response to α-Syn OE than previously recognized: at ∼5 weeks post-injection. Previous work from other groups have also demonstrated that there are conditions which cause increased spine density in response to α-Syn OE. One study investigated human α-Syn OE in newly born dentate gyrus cells of the hippocampus under both the conditions of a transgenic mouse line (PDGF h-α-Syn) or viral transduction of an overexpression construct (Winner et al., 2012). In the transgenic line, newly born neurons that overexpressed α-Syn within surrounding α-Syn OE tissue contained fewer dendritic branches, but higher spine density compared with newly born neurons in the non-transgenic line. Interestingly, individual neuronal transduction of α-Syn OE within wild type tissue led to reduced dendritic spine density (Winner et al., 2012). Although our current study did not investigate dendrite arborization, these published data suggest that both the local microenvironment and influence of neighboring neurons, as well as α-Syn expression levels within neurons can influence spine density, highlighting autonomous vs. non-cell autonomous differences (Winner et al., 2012).

Our confocal analysis provides context regarding the microenvironment. Cells in the region positive for pSer129 α-Syn did not show changes in dendritic spine density compared with those on the contralateral hemisphere. By contrast, neurons within the transduced region but negative for pS129 α-Syn demonstrated a relative increase compared with neighboring and contralateral cells. One limitation was that while we were able to stain pSer129α-Syn within individual neurons used for spine quantification, it was not possible to stain total (non-phosphorylated) synuclein at cellular resolution within the injected region (despite good staining elsewhere); we suspect that high α-Syn OE within the localized region prevents full penetration of the antibody through the tissue. Thus, it is possible that neurons with increased density have mildly elevated α-Syn levels that are below the level necessary to provoke pathological phosphorylation. Despite this limitation, our experiment demonstrates that the localized, regional overexpression of α-Syn induces increased spine density in cells without phosphorylated α-Syn, while cells with phosphorylated α-Syn show no difference from control cells. Given the large number of synuclein-positive axons in the region of the transfection site, it is also likely that presynaptic terminals from pSer129-positive neurons interact with spines from the pSer129-negative cells. One potential explanation for our results is that pre-synaptic α-Syn may have a stabilization effect on new spines, thereby increasing spine density. Interestingly and in contrast to previous studies, we did not detect any decreases in spine density in the dendrites that were pSer129 positive. Whether this is a feature of differential vulnerability of cortical neurons compared to other cell types, or due to the overexpression time (6 weeks) used in the confocal study, will be intriguing to investigate in future studies.

Our findings are contrasted by multiple studies in the literature showing decreased spine density following α-Syn perturbations. For example, studies in the olfactory bulb (Neuner et al., 2014) and cortex (Blumenstock et al., 2017) of transgenic synuclein overexpression mice demonstrated reduced dendritic spine density. Adult born granule cells of the olfactory bulb in A30P mice had reduced spine density compared with controls (Neuner et al., 2014). PDGF-h-α-Syn (α-Syn OE) mice have reduced cortical dendritic spine density starting at 3 months of age and persisting from 6 months onwards, along with greater turnover of spines in the OE group (Blumenstock et al., 2017). Interestingly, the decline in spine density did not get progressively worse after 6 months of age and no differences were seen between control and α-Syn OE mice in an earlier timepoint of 2 months (Blumenstock et al., 2017). Two major possibilities exist for the differences between our study and other models; 1) local overexpression presents neurons with different conditions within the context of an otherwise wildtype set of afferents from more distant brain regions. In other words, spine loss in other models may be due to dopaminergic or cholinergic depletion. 2) Our early timepoint represents a pre-symptomatic stage of α-Syn pathology where spine density is initially elevated. Future work to extend the experimental timeframe of imaging will be useful to delineate any potential ‘crossover’ point between increased spine density early and decreased density in response to chronic α-Syn OE.

The consequences of increased spine survival from our manipulation are not yet known. Experiences alter cortical dendritic spine anatomy and density (Nava et al., 2017, Pan et al., 2010); therefore, increased spine density can be advantageous under certain conditions. For instance, learning a new motor task leads to clustering of new spines near established spines that are related to a given task, and these spines are strengthened as the task familiarity increases in the primary motor (M1) cortex (Hedrick et al., 2022). Whether the increased spine density that we observe in response to α-Syn OE translates to measurable effects on PFC-mediated behavioral performance remains to be tested. Conversely, an aberrant increase in spine density may also lead to detrimental effects on brain networks. Davidson et al. (2020) recently found that spine density naturally increases with age in mouse Layer V neurons of M1 cortex (Davidson et al., 2020). In some studies, hyperexcitability coincides with increased spine density (Chen et al., 2012) and may be detrimental long-term to neuronal circuitry. Intra-parenchymal AAV injection of human α-Syn into rat neonates caused robust cortical overexpression (amongst other brain regions) and resulted in increased open field activity at 6 months of age, impaired neuronal health, hallmarks of apoptosis and reduced cortical ChAT immunopositivity (Aldrin-Kirk et al., 2014). The early timeframe over which we measured the effects of α-Syn OE limits our ability to conclude that local overexpression is not ultimately detrimental to cortical neurons, yet provides important insight into early responses to overexpression. Advances in techniques for extended longitudinal tracking using prisms will allow future studies to determine if cortical neurons remain less vulnerable over the lifetime of the animal and evaluate the behavioral consequences longitudinally. These data will be important for identifying why some neurons are especially vulnerable as an essential question for understanding Lewy Body Diseases.

### Limitations

One limitation of our study is that spines less than 0.45 µm (length) were not counted as they could not be distinguished from noise in vivo. Thus, an alternative explanation for the findings is that α-Syn increased the relative size, rather than number, of dendritic spines, leading to greater detection. This limitation is mitigated in part by our confocal study that allowed improved detection of smaller spines. Within the confocal study, spine density remained higher, and pSer129-negative neurons on the AAV-α-Syn injected side were on average longer, but not wider than spines from the contralateral side (Supplemental Fig 3).

Secondly, individual dendrites within the in vivo study had missing timepoints due to a variety of factors, including loss of window transparency and covid-19 pandemic related disruptions. For this reason, the primary outcomes were analyzed by independent statisticians to determine whether the conclusions were robust.

Finally, a major highlight of our study, distinct from prior experiments, was imaging of the same dendrites both before and after local viral overexpression, a powerful but challenging technique that allows evaluation of individual dendrites response to manipulation. However, this technique introduced two confounds described in the manuscript. First, introduction of the glass pipet directly at the site of imaging occasionally caused disruption or surface bleeding that may have impacted neuron health, spine density, and caused increased likelihood of premature opacity of the window. This limitation is likely to be equal between groups. Future use of a conditional, drug-driven strategy could mitigate this issue, but also has its own caveats as the drug could alter dendritic spines. Secondly, as discussed extensively above, expression of mCherry by AAV (tested with two different promoters) reduced visualization of genetically encoded, Thy1-driven YFP and GFP. This finding significantly limited our ability to separately evaluate transduced and non-transduced neurons. For this reason, we focused on mCherry-negative cells exposed to the local micro-environment to avoid this confound. Despite these limitations, this study shows a consistent response to early, local cortical overexpression and provides evidence that these changes are secondary to changes in spine survival.

### Conclusions

Lewy body diseases arise due to a complex interplay of genetic and protein perturbations that affect neuronal circuitry, which are then compounded by the long timeframe from prodromal to symptom onset. Evaluating the role of cortical α-Syn allows a better understanding of this complexity in this multi-brain-region, multi-system disease. The long-term goal is to understand the relative balance and interconnection of pathology in diverse brain regions, encompassing the heterogeneity from Parkinson’s disease with motor symptoms to pure-psychiatric onset disease. These insights are key to devising effective therapies to treat diverse Lewy-body related symptoms.

## Methods

### Animals and treatments

All experiments involving mice were approved by the University of Iowa Institutional Animal Care and Use Committee and were performed in strict adherence with the National Institute of Health (NIH) Guide for the Care and Use of Laboratory Animals. Mice were handled until comfortable with experimenter handling and with the Neurotar setup before any imaging sessions commenced.

For in vivo spine analysis studies, we utilized the *Thy1-YFP* mouse line, encoding the fluorescent molecule, YFP under the control of the Thy1 promoter: B6.Cg-Tg(*Thy1-YFP*)HJrs/J; stock number 3782 (Jackson Labs), and the *Thy1-GFP* line for confocal imaging studies: Tg(*Thy1-GFP*)MJrs/J; stock number 007788 (Jackson Labs). For *Thy1-YFP*, males and females were used and randomized to either control (n=6) or α-Synuclein overexpression (n=8) treatment groups prior to experiment onset. All mice were injected as mature adults, with age range for controls 39 – 70 weeks old; α-Synuclein overexpression range 40 – 73 weeks old. The *Thy1-GFP* mice contained 4 mice (3m, 1F); age at injection ranged from 11-24 weeks old.

### Cranial Window surgery

Cranial window implantation was performed as in our prior studies (Keyes et al., 2021) with changes as noted. Cranial window surgeries in *Thy1-YFP* mice were performed using ketamine/xylazine for anesthesia. Under deep anesthesia, animals were administered Meloxicam (2 mg/kg, IP) and Bupivacaine (8 mg/kg, SQ) prior to shaving and cleaning of the surgical site with Betadine surgical scrub and 70% ethanol. The skull was exposed using a clean incision from anterior-posterior with surgical scissors, periosteum was removed, and a 3 mm diameter round craniotomy was made using a dental drill with a 0.5 mm bit (Fine Science Tools). Cranial windows were installed covering a region −0.5 mm bregma to 2.5 mm bregma (AP), 2.5mm on the left, and 0.5mm on the right. The craniotomy crossed the sagittal suture and included the future injection site location. The dura was removed in all animals at the time of cranial window implant over the imaging region, to allow penetration of a pulled glass pipette with limited disruption to the imaging location. The exposed brain was covered using Kwik-Sil silicon adhesive (WPI), and the 3 mm coverslip window was placed to cover the craniotomy before being sealed with cyanoacrylate. The coverslip contained a pre-drilled hole, which was later used to inject viral vector into the brain. Two skull screws (#000-120) were placed over occipital cortex, inserted only deep enough to securely hold the skull. A model 11 stainless-steel lightweight headplate (Neurotar Ltd, Finland) was attached securely to the skull using cyanoacrylate followed by Teets dental cement (Methyl methacrylate). Mice were given normal saline (0.9% NaCl, SQ) post-surgery and allowed to recover for 3 weeks prior to imaging onset.

### Brain injections

#### Thy1-YFP for 2-photon microscopy

For *Thy1-YFP* mice we used the following viral treatments: Control: AAV6-CAG-mCherry-WPRE (1.7×10^12 gc/mL, Vector Biolabs, unilateral); α-Syn overexpression: vector coding for human, wild-type α-Synuclein: AAV6-CAG-hSNCA-WPRE (1.0×10^13^gc/mL, VectorBiolabs) was mixed with this mCherry virus outlined above. The ratio of synuclein to mCherry in the treatment group was adjusted from 3:1 to 1:1 following discovery of interference between mCherry and YFP fluorescence (n=3 synuclein mice). Animals were prepared for trans-window injection by cleaning the window and silicon plug as described for surgery. Anesthesia was induced using Isoflurane anesthesia using the Somnosuite® low flow anesthesia system (Kent Scientific). For *Thy1-YFP* mice, injections were done within 24 hours following imaging at “week 0” (**Fig. 2A**). Injections were made through the pre-existing hole in the glass coverslip through a silicon seal that was located approximately 1.25 mm lateral to midline (**Fig. 1A**). A pulled glass pipet, pre-loaded with virus, was directed with a 20 to 40-degree angle into the brain depending on the location of the hole relative to the target imaging region. 100-200nl of virus was injected at 5-6 locations along a track under the window, at approximately 0.4 mm, 0.8 mm, 1.2 mm, 1.6 mm, and 2 mm below the surface of the brain (maximum 1.2 µL total). Variations in amount given reflect adjustments made to better target cells in the imaged region, but were matched between groups.

#### Thy1-GFP for confocal microscopy

The *Thy1-GFP* mice (n = 4) received bilateral injections with control virus into one hemisphere and α-Synuclein overexpression into the other hemisphere: Control: AAV6-CAG-mCherry-WPRE (1.7×10^12^ gc/mL, Vector Biolabs, 1 μL, left hemisphere) and overexpression: vector coding for human, wild-type α-Synuclein: AAV6-CAG-hSNCA-IRES-mCherry-WPRE (1.0×10^12^ gc/mL, VectorBiolabs, 1 μL right hemisphere), which contains mCherry after an internal ribosomal entry site (IRES) to express mCherry in all α-Syn overexpression cells. For *Thy1-GFP* mice, we injected straight down (no angle). After the skull was exposed, bilateral craniotomies were drilled using a 0.5 mm drill bit at the following locations from bregma: 0.75 mm medial-lateral (ML), +/-1.4 mm anterior-posterior (AP), and −1.0 mm DV. As with the in vivo experiment, virus (total 1.0 μL/ hemisphere) was injected using gentle air pulses through a solenoid attached to an air supply through a pulled glass pipet at an approximate rate of 100nl/min. A greater proportion of virus (∼600 nL) was aimed at Layer V and the remaining (∼400 nL) was targeted to Layer II/III along the injection track. Following the completion of injection, the needle was left in place for 5 mins prior to withdrawal. After injection, the skin was sutured, normal saline (0.9% NaCl, SQ) was administered, and the mouse was allowed to recover for 6 weeks prior to trans-cardial perfusion and brain collection.

### 2-photon imaging

Mice were introduced to the Neurotar (Neurotar Ltd) air table 2 weeks prior to the first imaging session, and imaging began approximately 3 weeks after the cranial window surgery. Neurotar® is a tracking system and air table set up that allows animals to run on a free-floating carbon cage whilst head-fixed using a surgically implanted headplate. An overview dendritic map of the region was generated the week prior to the first spine imaging session. Imaging of YFP and mCherry were done using the following conditions: 935 nm laser wavelength, 3% offset, 0.5 mm step size, 320×320 pixels (0.1989 µm/pixel), 8x digital zoom. Voltage of the laser was adjusted dynamically for depth throughout the z-stack, as well as overall window clarity. The 25x objective correction collar was set at 0.17 for water, and 0.23 for aqueous jelly (Aquasonic clear ultrasound gel, Parker Laboratories Inc, diluted 4x in water). Backfill setting were adjusted for the highest resolution. Imaging sessions lasted up to 1 hour using an upright Olympus multiphoton FVMPE-RS equipped with Mai Tai DeepSee tunable laser set at 935 nm and a 25x water-immersion objective (N.A. = 1.05, Olympus). Mice were imaged each week for which chosen dendrites were clearly visible, up to 11 weeks consecutive weeks. In several animals, imaging was delayed beyond two weeks after window surgery to allow further clearing of the window.

### Immunostaining

Six weeks after injection, *Thy1-GFP* animals were heavily anaesthetized using ketamine/xylazine and trans-cardial perfusion was performed using cold 4% paraformaldehyde (PFA). Brains were removed, post-fixed overnight in PFA and cryoprotected using 30% sucrose in PBS. Brains were frozen in OCT and cryosectioned at 100 µm thickness to maximize the dendritic arbor that could be imaged. For immunostaining procedures, sections were blocked (2% Normal Goat serum, 0.3% Triton X-100 in PBS) and incubated with anti-GFP (chicken, Invitrogen, A10262, 1:100) and anti-phospho-synuclein (pSer129 rabbit, Abcam, ab51253, 1:500) primary antibodies diluted in blocking solution for 24 hours at 4°C with gentle agitation. Sections were washed with PBS and incubated with the relevant secondary antibody conjugated to Alexa-Fluor 488 nm (anti-chicken, A-11039, Invitrogen) or 647 nm (anti-rabbit, ab150079, Abcam) overnight in blocking solution at 4°C with agitation. They were then washed with PBS and mounted using Prolong™ glass Antifade Mountant (ThermoFisher) and cured for 48-60 hours prior to imaging. Confocal imaging was performed using a Leica TCS Sp5 microscope and images were collected using LAS-AF software. Confocal images were taken with 0.33 µm step size, using the appropriate collection methods for the relevant Alexa-Fluor molecules used in immunostaining. Images were analyzed using Imaris (see below) and adjusted in FIJI (Schindelin et al., 2012) or Adobe photoshop only for presentation.

### Analysis

#### 2-photon imaging analysis

Two photon imaging analysis was performed using FIJI. Dendrites and counting bins were selected based on predetermined criteria. Bins started at least 10 µm below the pial surface, and at least 10 µm away from any dendrite termination (potential growth cone). Bins were selected for analysis based on a target bin length of 15-25 µm and a minimum of 3 spines pre-injection. For analysis, two individuals blinded to the treatment group identified spines in all consecutive imaging weeks. Spines were counted if they were at least 0.45 µm long (>2 pixels). Spines that unambiguously disappeared for one week, then reappeared in a similar position the following week were recorded as new spines. To monitor transfection in the region of interest, the transfection of the dendrite was recorded, as well as the number of nearby transfected neurites within the imaging location of the dendrite (∼60 µm x 60 µm). Dendrites were defined as transfected if mCherry was visible in the dendrite when imaged using laser excitation of 1040 nm. Regions that contained local transfected neurites were scored as 0 (for no transfected neurites visible), low (for 1-5 neurites visible), or high (for 6 or greater neurites visible in the region).

For spine survival analyses, only spines that we recorded as ‘new’ during week 1-3 that were then lost during the imaging period were included. The number of weeks were counted from the week a spine appeared until the week it was recorded lost and then survival curves were generated. Analysis scripts were custom written using R, using the tidyverse package and libraries within. Kaplan-Meier curves and analysis were generated using ggsurv() and ggsurvplot() in the survminer package.

#### Confocal imaging analysis

Images that were collected on the Leica Sp5 microscope were collected as .lif files and imported directly into Imaris Microscopy Image analysis software (Oxford Instruments). Filaments were traced along basilar dendrites beginning at least 20 µm from the soma by an experimenter blinded to treatment conditions. Filaments which represented dendrites were used to automatically trace spines in Imaris and each spine was manually checked for accuracy and corrected if required. Filaments were not traced within 5 µm of a branchpoint or 10 µm from the dendrite terminal. All relevant data was exported in .csv format file and relevant data was extracted using custom R scripts and/or excel pivot tables. GraphPad Prism was then used to analyze the data, divided into 3 groups based on treatment and transfection status of the dendrite: control hemisphere/mCherry-negative; α-Syn OE hemisphere/pSer129-positive; α-Syn OE hemisphere/pSer129-negative. Full cell reconstruction from the confocal images were made using the FIJI plugin, “Pairwise stitching”, to stitch images together and the measure function was used in FIJI for the distance from soma to the center of the analyzed filament bin.

#### Statistics

Statistics were completed using R version 4.3.1, (for density and survival) and GraphPad Prism 9 (for turnover, persistence, morphology), and cross-checked by separate investigators. Data analysis for Figure 2C (spine density) and Figure 3C was performed by the Biostatistics core at the University of Iowa Institute for Clinical and Translational Science. For the analyses presented in Figure 2C, all follow-up spine densities were divided by that dendrite’s baseline density to obtain a measure of proportional change over time. Then a linear mixed effect model was fit to assess the effect of time, treatment, and the interaction between these two variables on this proportional change in density measure. A random intercept was used for each dendrite to account for inherent between-dendrite variability. The analyses presented in Figure 3C used a logistic mixed effects model in which the odds of a spine being persistent were modeled with time and treatment group serving as the predictor variables. Once again, a random intercept was used for each dendrite to account for inherent between-dendrite variability. Additional details and justification for other statistical tests are provided within the results section for clarity. P-values less than 0.05 were considered statistically significant for all analyses.

## Acknowledgements

Aly Yokimishyn, Sheyna Nathwani, Lucia Wagner, Aimee Bertolli, Patrick Ten Eyck, Yuriy M Usachev

---

**Supplemental Figure 1:**
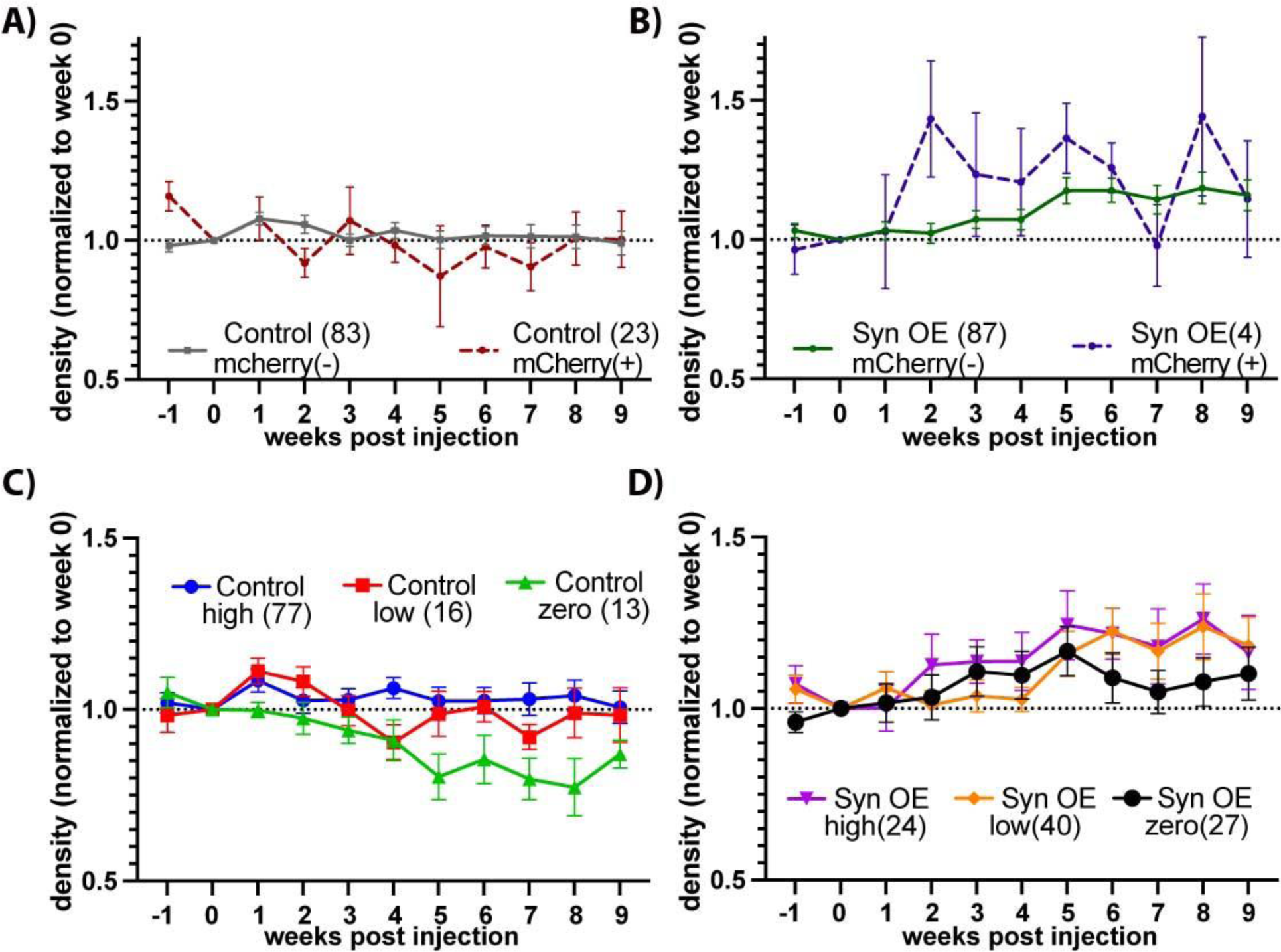
Effect of Transduction. **(A, B)**: Relative change in spine density based on detection (-/+) of mCherry at 1040 nm during live imaging, in mice injected with virus coding for mCherry (aka Control, A), or synuclein overexpression (Syn OE: mixed virus mCherry + α-Synuclein, B). Few mCherry positive dendrites could be reliably counted because high levels of mCherry caused loss of YFP, and low levels could not be detected by 2-photon at 1040nm. Dendrites with ambiguous transduction at 1040nm were excluded from this graph. Total identified dendrites are listed in parathesis but were not available for all weeks. **Local Microenvironment (C, D)**: As α-Synuclein may be secreted or impact presynaptic partners, we graphed how α-Synuclein in the layer I microenvironment affected dendrites localized to this area in mCherry injected mice (C) and in SynOE mice (D). Sub-analyses for illustrative purposes only; no statistics were performed given the small sample size within groups/weeks.

**Supplemental Figure 2:**
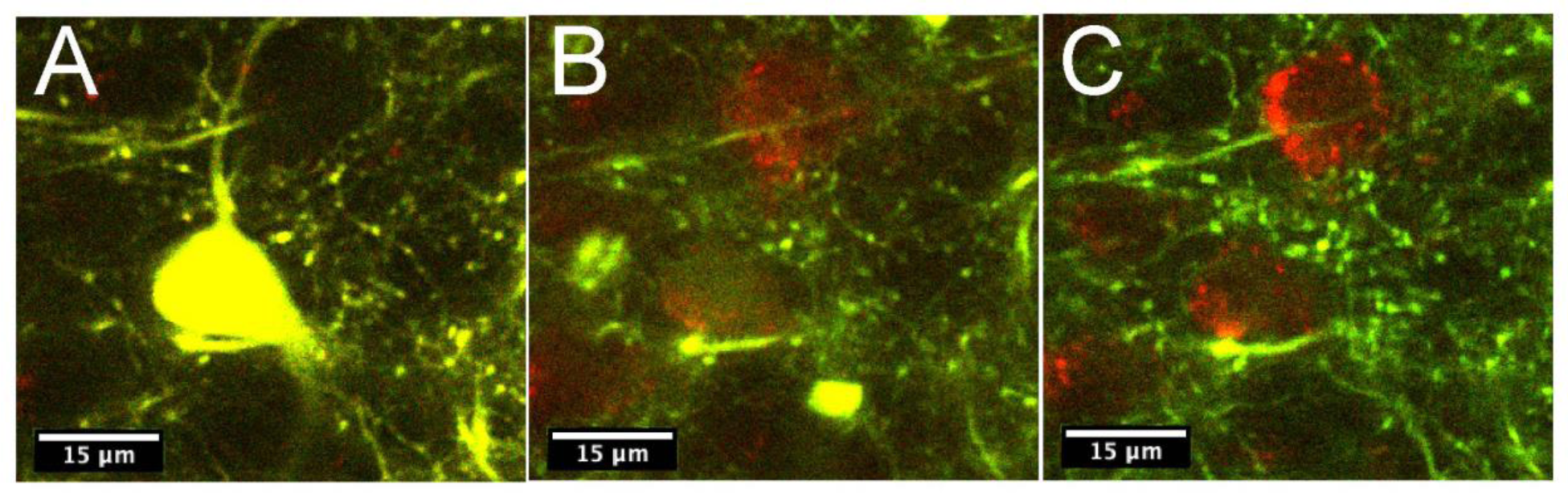
Detection of Yellow-fluorescent protein (YFP) expression in the Thy-1 YFP-H line decreases following transfection with AAV6 coding for mCherry. (A) pre-injection cell body, (B) 8 weeks following injection the YFP is barely visible in the cell body and mCherry is present, and (C) 11 weeks following injection YFP is no longer detectable and mCherry is clear. (Imaged at 935nm, which favors YFP). Similar issues were found with different transgenic models (GFP-M line), different serotypes (AAV1) and different promoters (hSYN vs. CAG). This phenomenon is mitigated when mCherry is preceded by IRES to reduce expression, as in our confocal experiment (See text), but this lower expression level is difficult to visualize in vivo.

**Supplemental Figure 3:**
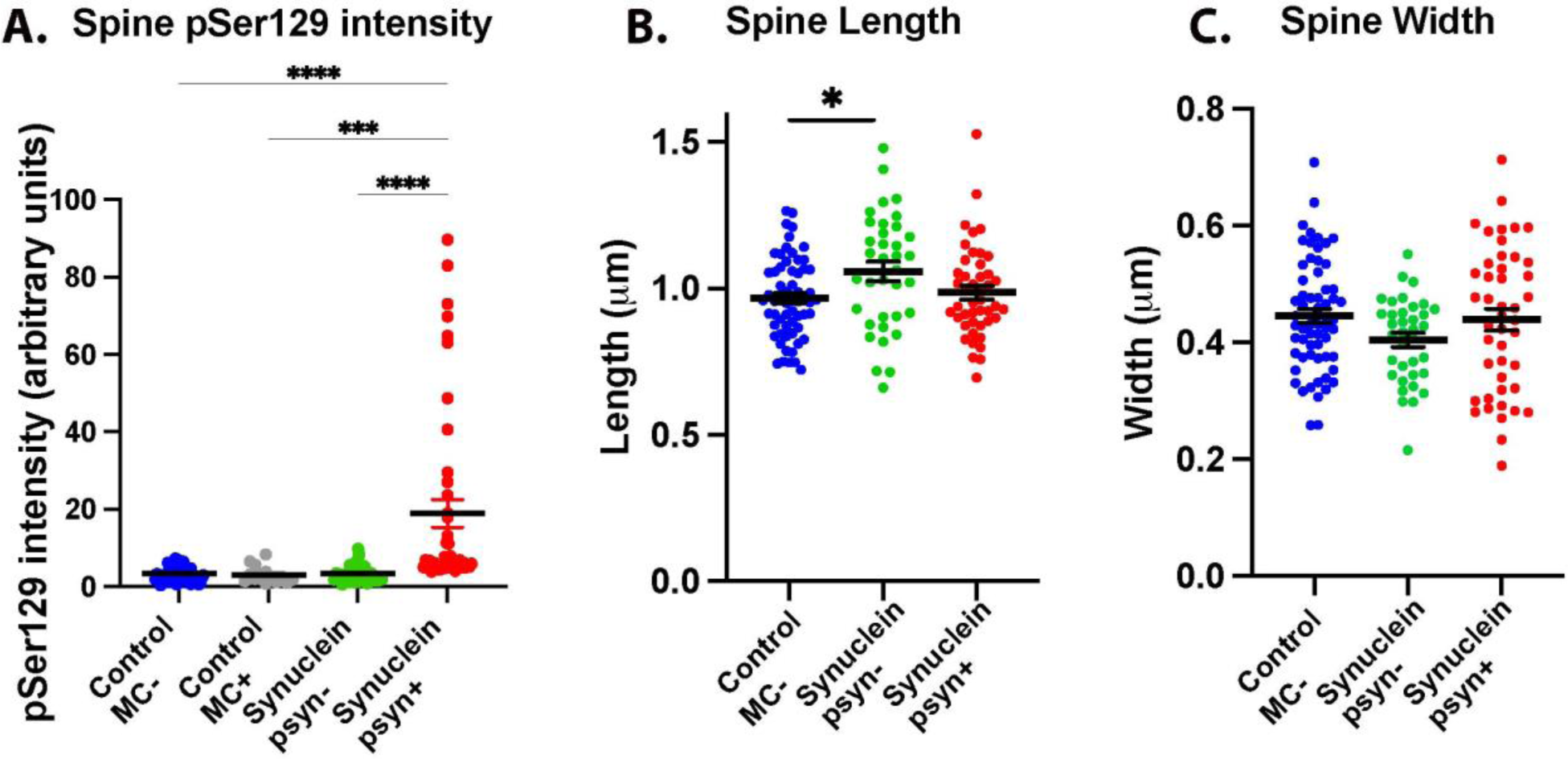
A) Phosphorylated α-Synuclein is detected in dendritic spines of dendrites that are pSer129 positive: Average phosphorylated α-Synuclein levels in spines from dendrites on the control side (mCherry negative and positive) and on the α-Synuclein side (pSer129 positive and negative). B) Spine length is longer in pSer129 negative dendrites on the α-Synuclein side compared with mCherry negative dendrites on the mCherry injected side. C) No significant differences were detected in spine width. mCherry positive dendrites on the mCherry injected-side were not analyzed due to reduced GFP (see limitations for details).

